# Treatment of anti-myeloperoxidase glomerulonephritis using recombinant deoxyribonuclease I is enhanced by adeno-associated virus gene therapy

**DOI:** 10.1101/2024.09.09.612148

**Authors:** Anne Cao Le, Virginie Oudin, Jonathan Dick, Maliha A. Alikhan, Daniel Koo Yuk Cheong, Mawj Mandwie, Ian E Alexander, A. Richard Kitching, Poh-Yi Gan, Grant J Logan, Kim M O’Sullivan

**Affiliations:** Centre for Inflammatory Diseases, Monash University Department of Medicine, 246 Clayton Rd, Clayton, VIC 3168, Australia; Gene Therapy Research Unit, Children’s Medical Research Institute and Sydney Children’s Hospitals Network, University of Sydney, Westmead, New South Wales, Australia; Discipline of Child and Adolescent Health, University of Sydney, Westmead, Australia; Departments of Nephrology and Paediatric Nephrology, Monash Health, 246 Clayton Rd, Clayton VIC 3168, Australia

## Abstract

Extracellular DNA (ecDNA) released from injured and dying cells powerfully induces injurious inflammation. In this study we define the role of ecDNA in systemic vasculitis affecting the kidney, using human kidney biopsies and murine models of myeloperoxidase anti-neutrophil cytoplasmic antibody-associated glomerulonephritis (MPO-ANCA GN). Twice daily administration of intravenous DNase I (ivDNase I) in two models of anti-MPO GN reduced glomerular deposition of ecDNA, histological injury, leukocyte infiltration and NETosis. Comprehensive investigation into DNase I modes of action revealed that after exposure to MPO, DNase I reduced lymph node DC numbers and their activation status, resulting in decreased frequency of MPO-specific CD4 effector T cells (IFN-γ, and IL-17A producing), and reductions in dermal anti-MPO delayed type hypersensitivity responses. To overcome the translational obstacle of the short half-life of DNase I (<5 hours), we tested an adeno-associated viral vector encoding DNase I (vec-DNase I). The method of DNase I delivery was more effective, as in addition to the histological and anti-inflammatory changes described above, a single vector treatment also reduced circulating MPO-ANCA titers and albuminuria. These results indicate ecDNA is a potent driver of anti-MPO GN and that DNase I is a potential therapeutic that can be delivered using gene technology.

## Introduction

Kidney involvement in ANCA-associated vasculitis is common, often due to autoreactivity to myeloperoxidase which results in the development of glomerulonephritis (GN). If left untreated disease will progress to end stage renal disease requiring dialysis or a kidney transplant. Current treatments for MPO-ANCA-associated glomerulonephritis (MPO-ANCA GN) have reduced the number of deaths but have significant side effects, both in their detrimental effects on protective immunity and in their metabolic effects, leading to infections, malignancy and cardiovascular disease (1, 2). Understanding the underlying disease mechanisms of MPO-ANCA GN affords the opportunity to develop new therapeutics that target critical components of nephritogenic immune pathways and potentially offer safer more specific treatments.

MPO-ANCA GN has a characteristic pattern of glomerular injury with considerable cell death and necrosis (3–6). Cellular injury or cell death can release pro-inflammatory damage-associated molecular patterns (DAMPs), including nuclear release of DNA as extracellular DNA (ecDNA). Neutrophils which are the critical disease initiating cells in MPO-ANCA GN produce a unique form of cell death known as neutrophil extracellular traps (NETs) which expel large strands of DNA and is identified through de-condensation of chromatin. This process relies heavily on reactive oxygen species (ROS) and protein-arginine deiminase 4 (PAD4) dependent citrullination of histones (7–11). NETs are thought to occur to increase the bactericidal killing capacity of neutrophils by extending the surface area in which to entrap bacteria (7, 12). However, in the context of sterile inflammation NETs are highly injurious releasing over 70 known proinflammatory mediators. (13, 14). Fragments of ecDNA (including CpG oligonucleotides) are a major product of NETs as well as other forms of cell death (apoptosis, necroptosis, necrosis) and drives inflammatory gene expression. Of particular interest is detection of ecDNA and signaling through cytosolic and membrane-bound DNA sensors such as Toll like receptor 9 (TLR9), stimulator of interferon genes (STING) and cyclic GMP-AMP synthase (cGAS) as they induce inflammatory responses in sterile inflammation through NFkB nuclear translocation, type 1 interferon cytokine production (IFN-I) and inflammasome activation (11, 15–22). In host defense, excessive DNA is controlled by endogenous DNase enzymes (23). However, if cellular injury is severe, the production of ecDNA may exceed the capacity for DNase clearance thereby further driving inflammation.

The kidney is a major source of DNase I outside the digestive system and gene deletion of the enzyme in experimental mice induces GN (23–27). These observations support the possibility that DNase I removal of ecDNA is an important physiological process in the renal system. In support of this possibility, sera from patients with acute MPO-ANCA GN have a significant reduction of DNase I and higher levels of serum ecDNA when compared with healthy donors (28). Neutrophil extracellular traps (NETs) occur at high frequency within the glomeruli of patients with MPO-ANCA GN and are a large source of ecDNA (16, 19, 29). Importantly, it has been observed that MPO, the autoantigen in this disease, exerts enhanced biological effects when tethered to the ecDNA in NETs (30). These findings support the notion that ecDNA released from injured and dying glomerular cells and NETs, can exceed the capacity for DNase I clearance resulting in ecDNA accumulation in acute disease. The findings also support the possibility that exogenous DNase I administration may restore homeostatic clearance of ecDNA to relieve pathogenic inflammation.

In the current study, we sought to confirm whether ecDNA accumulation occurs in MPO-ANCA GN. Kidneys from patients with acute MPO-ANCA GN exhibited increased ecDNA, as well as reduced renal DNase I expression levels compared with control kidneys. The functional relevance of these phenomena was demonstrated in two murine models of anti-MPO GN where disease is mediated by MPO-specific T cells or by anti-MPO antibodies. Importantly, intravenous administration of exogenous DNase I (ivDNase I) in two different experimental models (cell mediated and MPO-ANCA mediated) markedly attenuated the development of GN indicating that DNase I is a potential treatment for MPO-ANCA GN. To overcome the translational obstacle of the short half-life of DNase I in serum (<5 hours), we explored the use of DNase I gene transfer using an adeno-associated viral vector (AAV, vec-DNase I) to convert the liver into a bio-factory for enzyme secretion into the circulatory system. The continuous supply of vec-DNase I in active murine anti-MPO-GN showed enhanced therapeutic efficacy over twice daily doses of ivDNase I, demonstrated by better preservation of glomerular histology, reductions in MPO-ANCA and albuminuria. Collectively these findings support the pursuit of DNase I for the treatment of MPO-ANCA GN through DNase I gene therapy.

## Results

### EcDNA accumulates in glomeruli of patients with MPO-ANCA vasculitis

To semi-quantitate the extent of ecDNA accumulation in kidneys of MPO-ANCA GN patients presenting with acute GN, we analyzed tissues collected from patients with this disease (*n*=29) and compared them with control tissues collected from patients with a non-proliferative form of GN, minimal change disease (MCD, *n*=6). The MPO-ANCA GN patients all had a positive serum MPO-ANCA assay and presented with severe renal disease with an estimated glomerular filtration rate (eGFR) of 35±8 (mean ±SEM, ml/min/1.73^2^) compared to MCD that had the expected normal renal function (eGFR 100±6). There was a mean of 18 glomeruli per biopsy (see other patient clinical data in Supplementary Table 1).

Renal biopsies were analyzed using a methodology that detects ecDNA regardless of the type of cell death (i.e. apoptotic, necrotic or NETotic). Fragmented ecDNA outside the nucleus of cells was assessed by semi-quantitative analysis of positive staining and expressed in arbitrary units per a previously validated method (31). There was significantly greater ecDNA deposition within the glomeruli of MPO-ANCA GN biopsies compared to control biopsies (Figures 1, A-C). It has been reported that patients with MPO-ANCA GN have lower levels of circulating DNase I (17), which may impair NETs digestion and removal of apoptotic bodies. A subset of the 29 renal biopsies used to measure ecDNA were labelled (*n*=6, the biopsies with sample remaining) with DNase I antibody demonstrated that MPO-ANCA GN patients have less DNase I expression within the tubulointerstitium and glomeruli when compared to those from control MCD patients (*n*=6) (Figure 1D and E).

**Figure 1.**
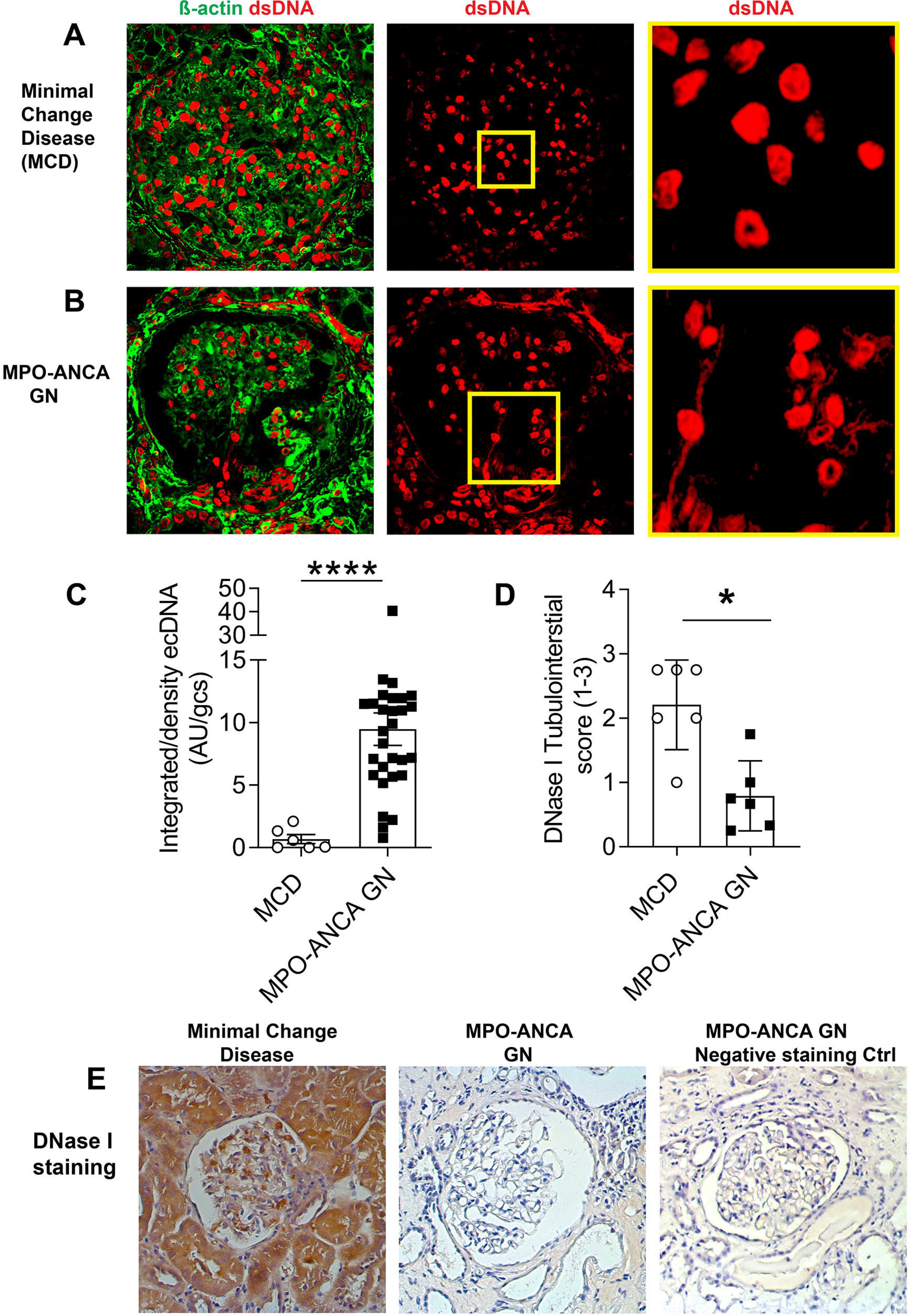
Enhanced extracellular DNA deposition and reduced DNase I expression in kidney biopsies from patients with MPO-ANCA GN. Renal biopsies from ANCA-vasculitis patients were stained for double-stranded DNA (dsDNA) (red) and counterstained with β-actin (green) to aid in identification of glomeruli and cells. (**A**) Biopsies from patients with minimal change disease (MCD) (with minimal glomerular injury) served as controls. Inset box shows higher power field of view with minimal ecDNA. (**B**) Patients with MPO-ANCA glomerulonephritis show extensive peri-glomerular and glomerular ecDNA. (**C**) Semi-quantitation of extracellular dsDNA deposition in MPO-ANCA GN patients. (**D**-**E**) MCD patients show extensive DNase I expression compared to the tubulo-interstitium of patients with MPO-ANCA GN. Negative control for antibody specificity confirms the significant diminution of DNase I in MPO-ANCA GN. *P<0.05, ****P<0.0001 Data are means ± SEM Human Data is from *n*=29 (MPO-ANCA GN) and *n*=6 (MCD) in each group and analyzed by Mann-Whitney U test. MPO, myeloperoxidase, ANCA anti neutrophil cytoplasmic antibody, GN, glomerulonephritis, Ctrl, Control; DNase I, Deoxyribonucleic I; dsDNA, double stranded deoxyribonucleic acid, ecDNA, extracellular deoxyribonucleic acid. Original magnification 400x, HP inset 3000x.

### Enhanced deposition of ecDNA and reduced expression of DNase I in the kidneys of mice with anti-MPO GN is attenuated with exogenous DNase I administration

Using an established murine model of anti-MPO GN (38), we tested the hypothesis that disease is associated with increased levels of ecDNA deposition in murine kidneys. Mice were immunized with MPO and GN was induced with anti-GBM globulin (Figure 2A, anti-GBM Ig). Glomerular injury in this model of active anti-MPO autoimmunity is mediated by cellular effectors and OVA-immunized mice injected with anti-GBM globulin serve as negative controls. Compared with control mice, mice with anti-MPO GN exhibited higher levels of ecDNA deposition and significantly less DNase I expression (Figure 2B). Thus, this model shows the same pattern of injury with respect to ecDNA and decreased DNase I expression as observed in the human disease (in previous Figure 1A). To test whether the reductions in endogenous DNase I levels could be offset using exogenous DNase I to alleviate clinical endpoints of disease, anti-MPO GN mice were given twice daily injections of human DNase I for four days. This treatment significantly attenuated glomerular ecDNA deposition (Figures 2B-C) and restored renal expression of DNase I when compared to control animals (Figure 2D). As expected, endogenous DNase I levels were unaffected in OVA-immunized control mice, which also did not develop anti-MPO autoimmunity as seen by the absence of GN, low levels of ecDNA deposition, NET formation and MPO deposition (Figure 2 B-E).

**Figure 2.**
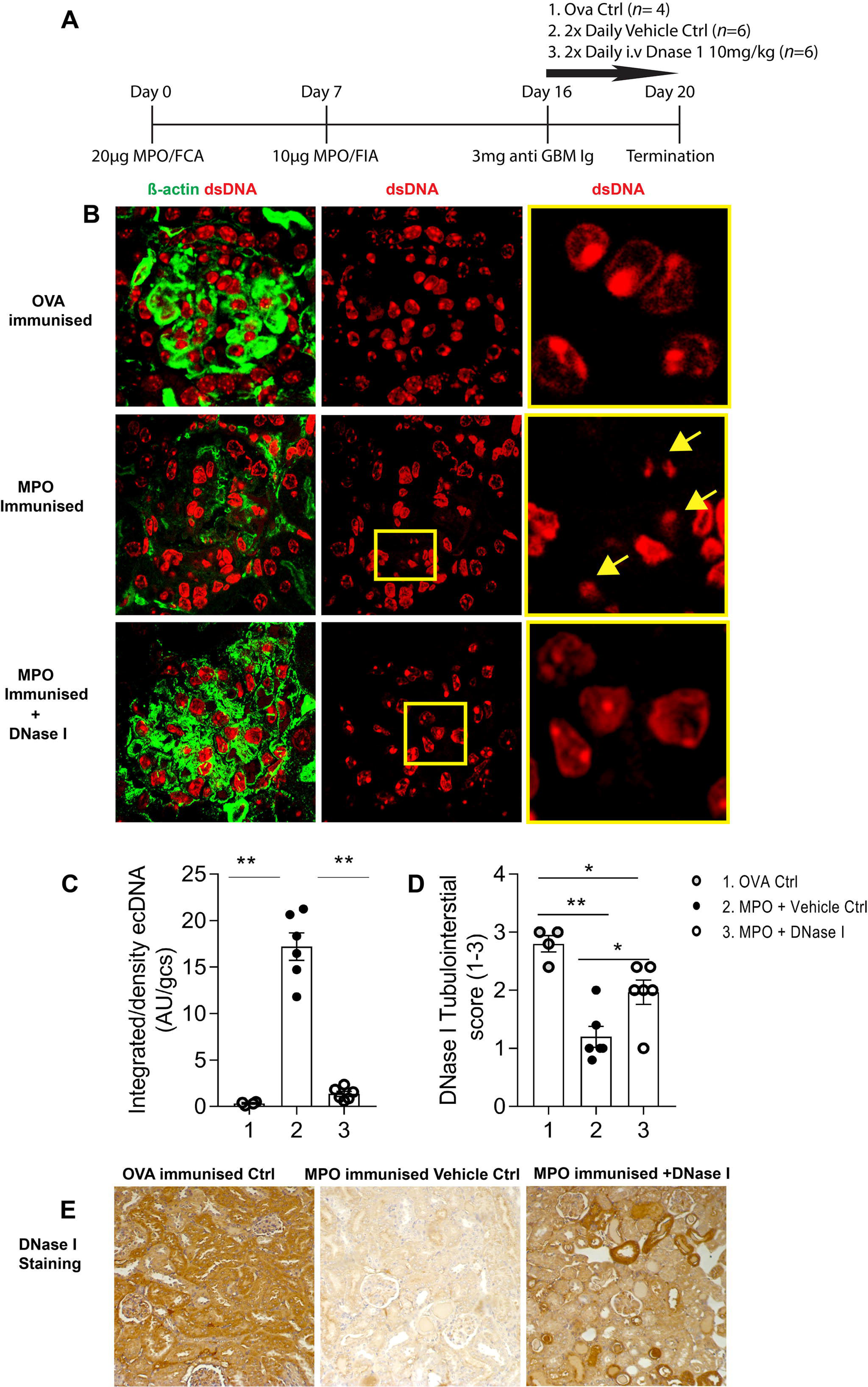
Mice with autoimmune anti-MPO GN have significantly enhanced renal deposition of ecDNA and diminution of DNase I, which is restored by ivDNase. **I.** (**A**) Experimental timeline outlining the induction of anti-MPO GN via immunization and sub nephritogenic dose of anti GBM Ig with : Ovalbumin (*n*=4) control (irrelevant antigen), MPO immunized and vehicle treated control (*n*=6) and MPO immunized, treated with DNase I (*n*=6). (**B**) Representative kidney sections from mice immunized with OVA, MPO-immunized vehicle treated mice or MPO-immunized mice that were later treated with ivDNase I. Sections were stained for ecDNA (red) and counter-stained with β actin (green). (**C**) Semi-quantitation of ecDNA in kidney sections made from OVA (1), MPO-immunized vehicle treated (2) and MPO-immunized ivDNase I-treated groups (3) was performed using Image J and the amount of ecDNA is expressed as arbitrary units per glomerular cross section (AU/gcs). (**D**) with Histological scoring of the tubulointerstitium of mouse kidneys for DNase I (x-axis is as panel C) (**E**) Representative images of DNase I immunohistochemistry across experimental groups. *P<0.05, **P<0.01. Data are mean ± SEM from six mice in each group analyzed by Mann-Whitney U or Kruskal-Wallis, for 3 or more groups. Abbreviations: MPO, myeloperoxidase, ANCA anti neutrophil cytoplasmic antibody, GN, glomerulonephritis, Ctrl, Control; DNase I, Deoxyribonuclease I; ecDNA, extracellular deoxyribonucleic acid; dsDNA, double stranded deoxyribonucleic acid; OVA, Ovalbumin. Original magnification 400x.

### Exogenous DNase I treatment diminishes glomerular injury and glomerular NET formation in MPO-ANCA GN

Having demonstrated that DNase I cleared ecDNA and maintained homeostasis of endogenous renal DNase I, further endpoints were assessed to determine the effect of enzyme administration on histological and functional injury and renal inflammation (Figure 3A, experimental design). DNase I treated mice exhibited less glomerular injury as seen by lower proportions of abnormal glomeruli (Figure 3B-C), with less incidence of glomerular crescents, segmental necrosis, glomerular expansion and cell infiltration (Supplementary Figure 1 A-D),less fibrin deposition (Figure 3D-E) and fewer glomerular leukocytes when compared to saline control mice that developed anti-MPO GN (Figure 3F-H). The mean albuminuria over 24 hours was numerically lower in DNase I treated mice but this did not reach statistical significance (P=0.06, Figure 3H). DNase I was administered to mice 16 days after establishment of autoimmunity and GN initiated with anti-GBM globulin. Despite only 4 days of treatment, DNase I treated mice had a significant reduction in dermal DTH, 24-hours after injecting the footpad with MPO, compared with the saline treated control group (Figure 3J). A significant decrease in MPO-specific IFN-γ and IL-17A producing cells from lymph node draining sites of MPO immunization was observed in the DNase I treated group (Figures 3K-L).

**Figure 3.**
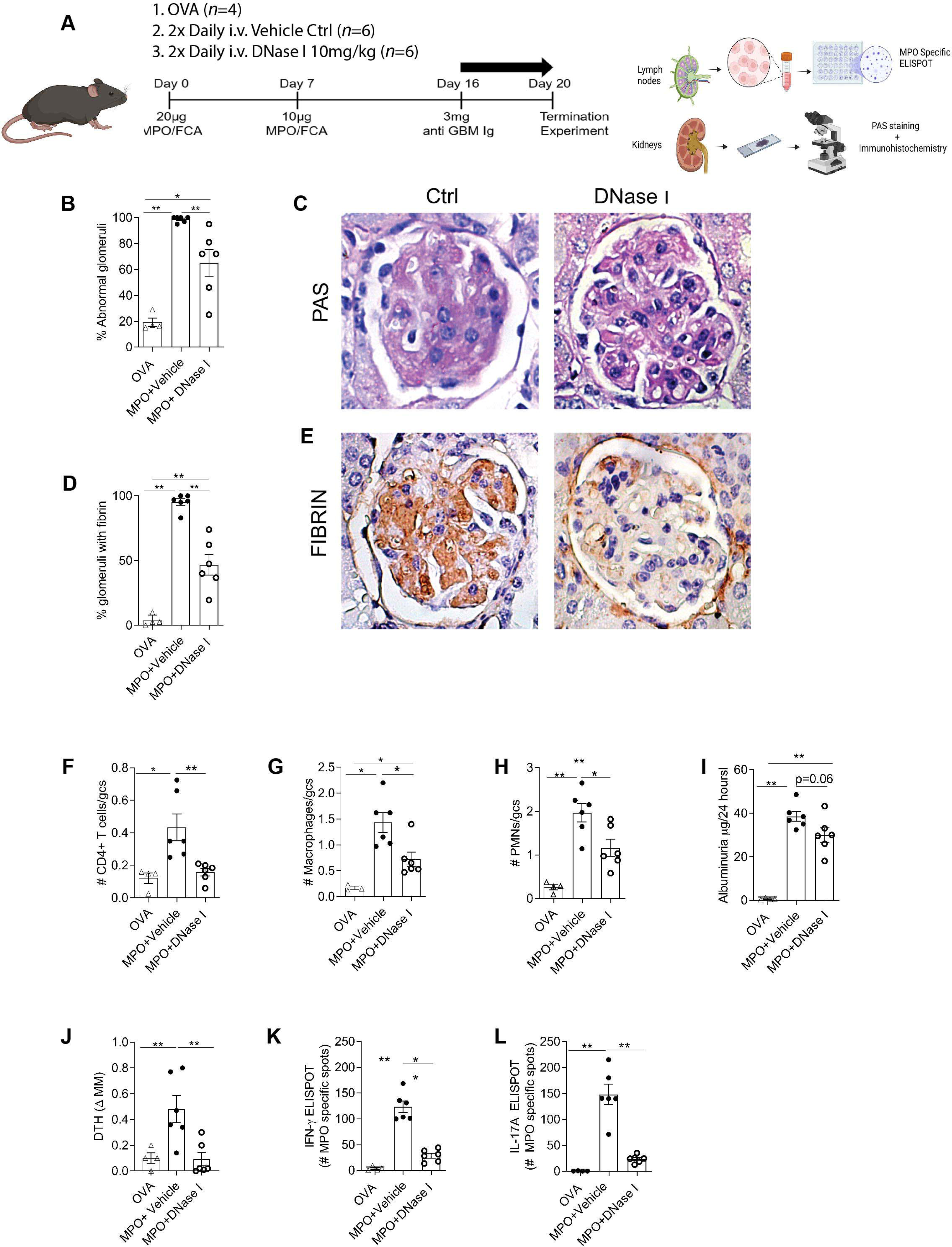
DNase I treatment reduces histological glomerular injury, leukocyte recruitment and functional injury in experimental MPO-ANCA GN. (**A**) Experimental outline and endpoints. Animals were assessed for the following parameters; (**B**) percentage of abnormal glomeruli after the assessment of (**C**) murine kidney sections stained with periodic acid-Schiff, (**D**) percentage of glomeruli containing fibrin after assessment of (**E**) murine kidney sections stained with an anti-fibrin antibody. Immunoperoxidase staining for (**F**) CD4+ T cells, (**G**) macrophages and (**H**) neutrophils. (**I**) Twenty-four-hour albuminuria. Animals were also assessed for (**J**) dermal delayed hypersensitivity response after intradermal MPO injection, (**K**) frequency of MPO-specific IFNγ and (**L**) frequency of MPO-specific IL-17A producing cells in lymph nodes draining sites of MPO immunization. *P<0.05, **P<0.01. Data are mean ± SEM from six mice in each group analyzed by Mann-Whitney U test. MPO, myeloperoxidase;DNase I, Deoxyribonuclease I; Ctrl, Control; DNase I, Deoxyribonuclease I; Original Magnification 600x.

We then sought evidence of the therapeutic efficacy of DNase I to reduce the number of NETs depositing MPO in the glomeruli of kidneys of mice with anti MPO-GN. In the control saline-treated mice NETs were found in 80% of glomeruli, while DNase I treatment markedly reduced the proportion of glomeruli with NETs to less than 10% (Figure 4A). This reduction in NETs was associated with less extracellular MPO deposition (Figure 4B-D).

**Figure 4.**
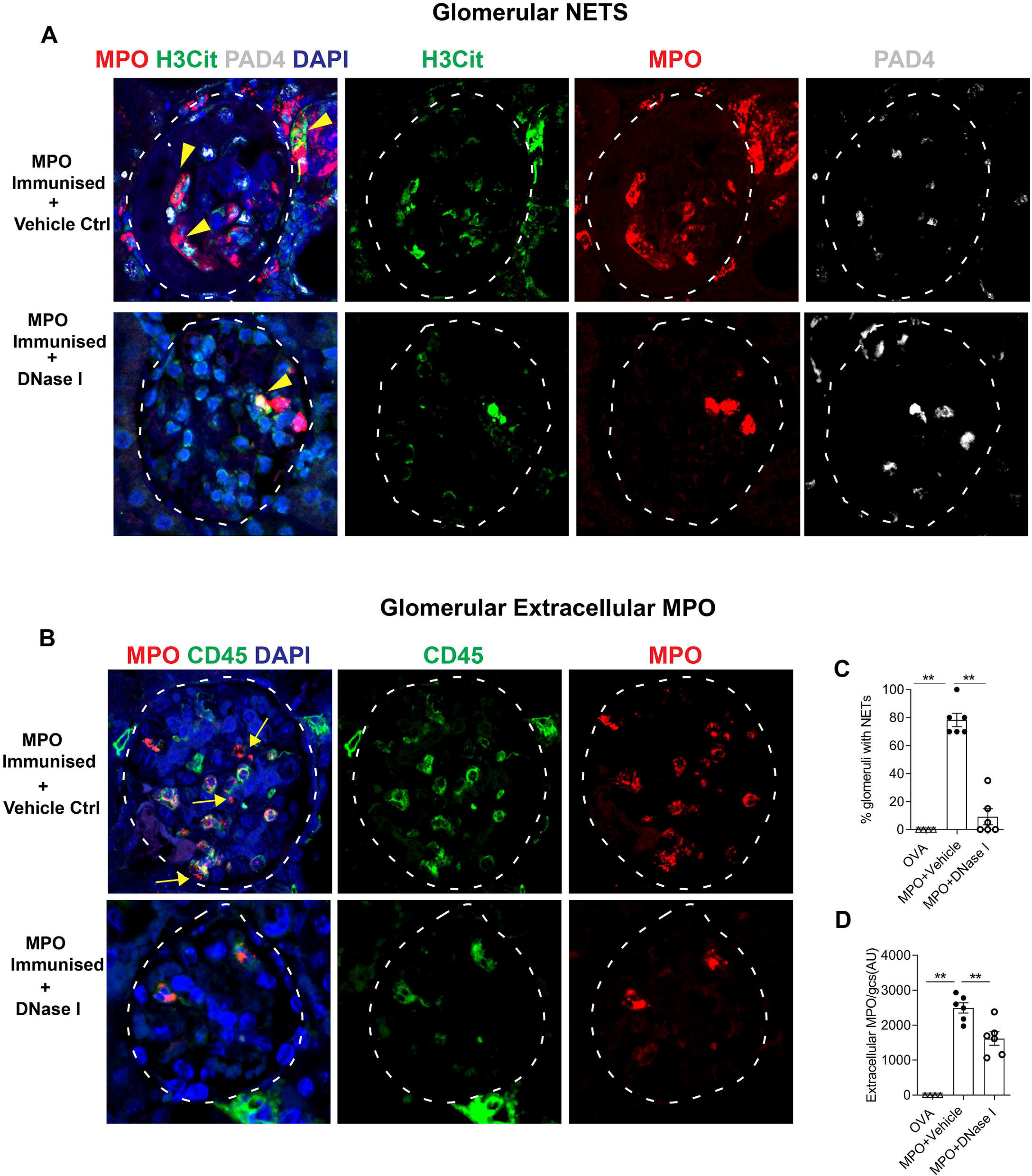
DNase I treatment reduces NET formation and deposition of extracellular MPO in established anti-MPO-ANCA GN. (**A**) Kidney sections from MPO-ANCA GN mice receiving either saline control or ivDNase I treatment stained for NETS by co-localization of MPO (red), H3Cit (green), PAD4 (white) and DAPI (blue), indicated by yellow arrow heads. (B) Kidney sections labelled for MPO (red), CD45 (green) and DAPI (blue) were assessed for intracellular MPO (co-localization of CD45 and MPO, arrow head) and extracellular MPO (non-co-localized CD45 and MPO, arrows). (**C**) NETS quantitation in glomeruli of ivDNase I treated and control MPO-ANCA GN mice. (**D**) Extracellular MPO quantitation in glomeruli of ivDNase I treated and control MPO-ANCA GN animals. **P<0.01. Data are mean ± SEM from six mice in each group analyzed by Mann-Whitney U. Anti MPO-ANCA GN, MPO, myeloperoxidase; DNase I, Deoxyribonuclease I; Ctrl, Control PAD4, peptidylarginine deiminase; H3Cit, Citrullinated Histone 3; DAPI, 4’, 6-diamidino-2-phenylindole, Ctrl, Control; DNase I, Deoxyribonuclease I; CD45, pan leukocyte marker C45. Original Magnification 600x

### Exogenous DNase I attenuates the development of anti-MPO autoimmunity

To assess the effect of exogenous DNase I on the development of anti-MPO autoimmunity, DNase I or vehicle control was administered to C57BL/6 WT mice (*n*=10 each group) one day prior to the groups receiving subcutaneous immunization with rmMPO in FCA, followed by daily injections of saline or DNase I for a further ten days (Figure 5A). When compared to saline control mice, DNase I treated mice had a significant reduction in dermal DTH to MPO (Figure 5B), but serum MPO-ANCA titers were unchanged (Figure 5C). *Ex vivo* MPO stimulation of cells from the lymph nodes that drain immunization sites demonstrated that DNase I reduced the numbers of IFN-γ secreting cells detected via ELISPOT (Figure 5D) whereas IL-17A production was unaltered (Figure 5E). Moreover, draining lymph node cells in this assay had a significant increase in CD4^+^Foxp3^+^CTV^-^ cells (Figure 5F). Collectively, this data strongly indicates that DNase I inhibits the development of anti-MPO autoimmunity by both reducing the frequency of Th1 MPO-specific effector cells whilst increasing the frequency of MPO-specific T regulatory cells with increased capacity to respond to MPO.

**Figure 5.**
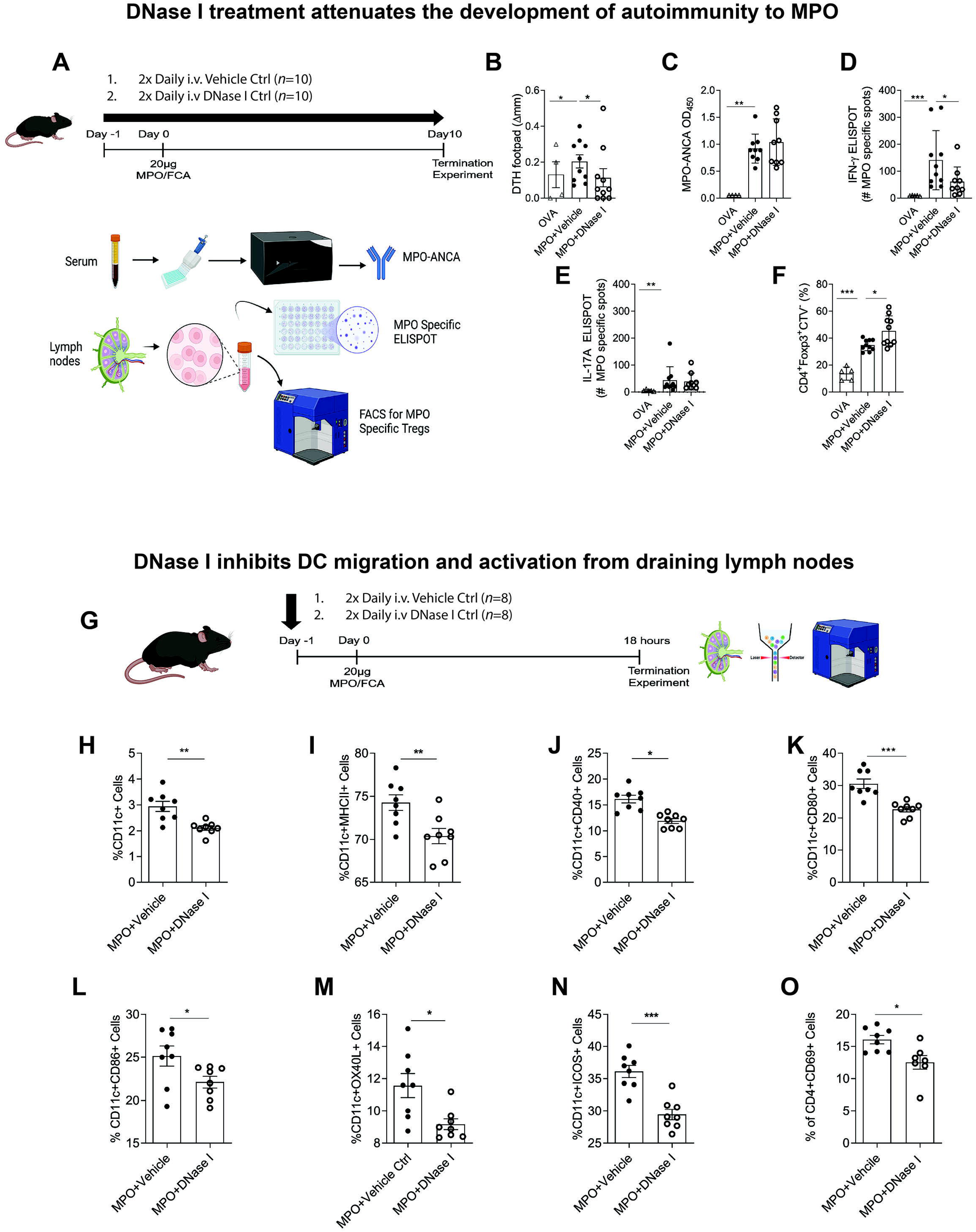
DNase I treatment reduces autoimmunity to MPO and reduces dendritic cells and T cell activation from the draining lymph nodes. (**A**) Experimental timeline for a 10-day MPO immunization model where mice receive saline of ivDNase I one day prior to immunization with MPO. Mice subsequently receive saline or ivDNase I daily until the end of the experiment at ten days post-immunization. Animals were assessed for (**B**) delayed type hypersensitivity response after intradermal MPO injection, (**C**) serum MPO-ANCA and frequency of (**D**) MPO-specific IFNγ, (**E**) MPO-specific IL-17A via ELISpots and (**F**) number of *ex vivo* CD4+FoxP3+CTV^-^ lymphocytes that proliferate in response to MPO, from lymph nodes from draining sites of MPO immunization. (**G**) Experimental timeline for an 18-hour model MPO immunization model where mice receive saline of ivDNase I one day prior to immunization with MPO with experiment terminated 18 hours post-immunization. Lymph nodes draining the site of MPO immunization were assessed for the frequency of (**H**) CD11c dendritic cells that also expressed (**I**) MHCII, (**J**) CD40, (**K**) CD80, (**L**) CD86, (**M**) OX40L or (**N**) ICOS. (O) Frequency of activated CD4 T cells in lymph nodes draining the site of MPO immunization. *P<0.05, **P<0.01, ***P<0.001, analyzed by Mann-Whitney U test. Data are mean ± SEM from n=10 mice in the 10-day model and n=8 mice in the 18-hour model. D-1, day minus 1; FCA, Freunds Complete adjuvant; µg, micrograms; kg, kilograms; IV, intravenous tail vein injection; DTH, delayed type hypersensitivity MPO, myeloperoxidase; IFNγ, interferon gamma; IL-17A, interleukin 17A, ELISpot, enzyme-linked immunoSpot; DNase I, deoxyribonuclease I; Ctrl, U. CD80, Cluster of differentiation; CD86, Cluster of differentiation 86; CD40, Cluster of differentiation 40 (co-stimulatory molecules); OX40L,; ICOS, Inducible T-cell co-Stimulator; MHCII, major histocompatibility complex; CD69, Cluster of differentiation 69 (marker of T cell antigen activation). DNase I, Deoxyribonuclease I; Ctrl, Control.

### Exogenous DNase I inhibits DC migration and activation in lymph nodes draining MPO immunization sites

To test the hypothesis that DNase I modulates DC activation after MPO immunization, the effects of DNase I were examined at the time of antigen presentation. C57BL/6 mice received ivDNase I 24-hours before immunization with rmMPO in FCA and experiments ended 18 hours later (Figure G). Draining lymph nodes were removed and CD11c+ cells examined by flow cytometric analysis for co-expression of activation markers. Eighteen hours after immunization with MPO, the mice treated with ivDNase I had a significant reduction in the proportion of CD11c+cells (Figure 5H). Also, DCs migrating to the draining lymph nodes showed a reduction in the expression of activation and co-stimulation markers MHC class II, CD40, CD80, CD86, OX40L, and ICOS, and (Figures 5, I-N). Mice treated with ivDNase I also had a significant reduction in the proportion of activated T cells (CD4+CD69+, Figure 5O). These findings are consistent with DNase I inhibition of DC activation and migration to draining nodes.

### DNase I attenuates disease in experimental GN induced by transfer of anti-MPO antibodies

As the International Society of Nephrology recommend therapeutic drug testing is undertaken in multiple pre-clinical animal models (32) we utilized a second animal model of anti MPO GN. As anti-MPO antibodies are important in activating neutrophils in MPO-ANCA GN, the therapeutic efficacy of DNase I shown in the 20-day T cell-mediated model of anti MPO-GN was also tested in a model of anti-MPO GN induced by transfer of anti-MPO antibodies into mice primed with LPS (33). DNase I (or vehicle) was given 24 hours prior to disease induction and 12 hourly until the end of the experiment (Figure 6A). Renal ecDNA deposition was significantly less (Figure 6B-C, *p*<0.05) and DNase I expression was significantly greater in ivDNase I treated mice than in vehicle-treated mice with GN (Figure 6D-E, *p*<0.05). Glomerular NET formation (Figure 7A) and deposition of extracellular MPO (Figure 7 B) were both significantly greater (*p*<0.05) in control vehicle-treated mice than in animals that received ivDNase I (Figure 7 C-D). Histological assessment of glomerular injury demonstrated ivDNase I treatment prevented the development of abnormal glomeruli and infiltrating immune cells (Figure 8 A-B). As in the 20-day active model, there was a trend to reduced albuminuria (Figure 8 C), a reduced number of abnormal glomeruli defined as a reduction in segmental necrosis and infiltration of cells in to Bowman’s space as described previously (Figure 8D) and reduced numbers of glomerular infiltrating neutrophils (Figure 8 E), and macrophages (Figure 8 F). Analysis of kidney mRNA for common inflammatory markers demonstrated significant diminution of chemokines that control macrophage recruitment (CCL2), (Figure 8G) neutrophil recruitment (CXCL1), (Figure 8 H) and markers of inflammation including IFNγ, CXCL2, IL-1β, IL-6 and TNF (Fig 9 I-M). Analysis of DNase I mRNA showed that non-treated diseased mice had significantly more intrarenal DNase I gene expression than the iv DNase I treated group, suggesting increased DNase I gene transcription in response to ecDNA accumulating in the kidney (Figure 9N). Collectively, the data from two different and complementary models of anti-MPO GN indicate that ecDNA deposition can be cleared by exogenous DNase I, resulting in significantly attenuated glomerulonephritis.

**Figure 6.**
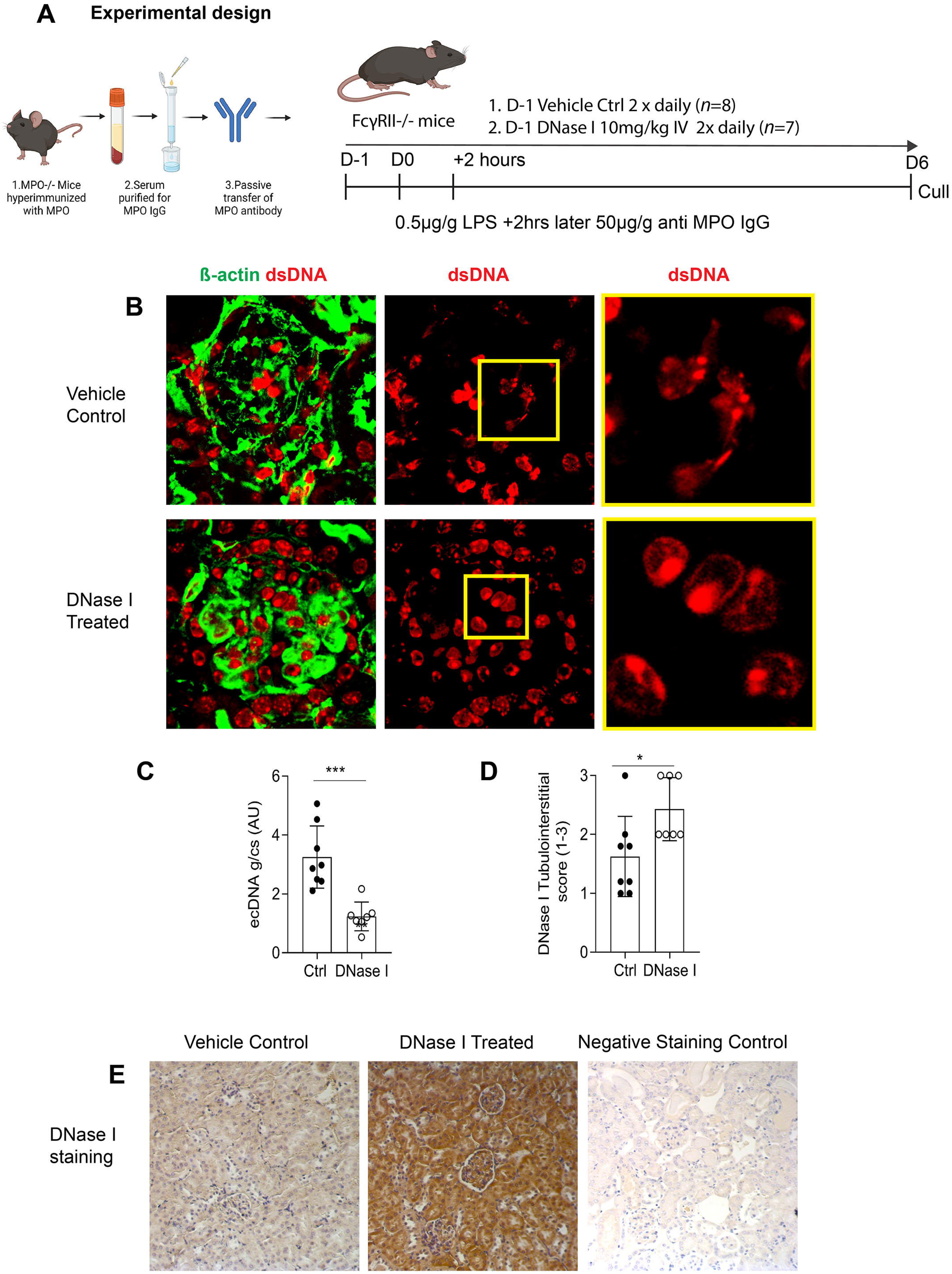
Extracellular DNA is reduced in DNase I-treated mice with GN induced by passive transfer of anti-MPO antibodies. (**A**) Experimental timeline, where mice receive saline or ivDNase I 1-day prior to receiving LPS and anti-MPO IgG (**B**) Kidney sections from mice receiving either saline control or ivDNase I treatment stained for DNA (red) and b-actin (green) were assessed were quantitatively assessed using Image J for (**C**) ecDNA (expressed in arbitrary units) per glomeruli cross-section. (**D**) Histological scoring of the tubulointerstitium of mouse kidneys for DNase I was performed on (**E**) kidney sections immunohistochemically-labelled for DNase I. *P<0.05, ***P<0.001. Data are mean ± SEM from 8 mice in the control group and 7 mice in the treatment group analyzed by Mann-Whitney U or Kruskal-Wallis for 3 or more groups. Abbreviations: Ctrl, Control; DNase I, Deoxyribonuclease I; ecDNA, extracellular deoxyribonucleic acid. Original magnification 400x

**Figure 7.**
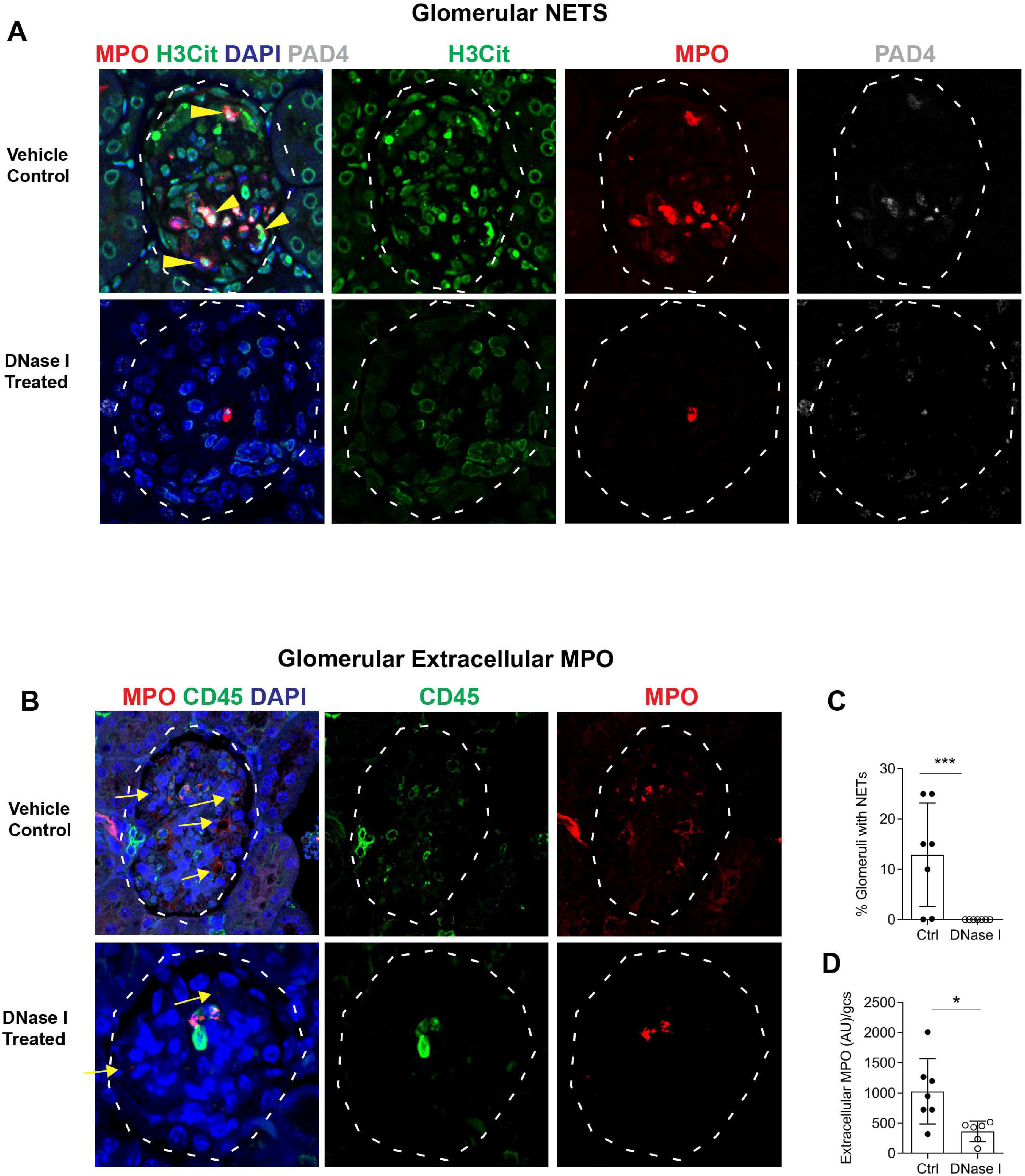
NET formation is abolished by DNase I treatment in a passive ANCA-dependent model of anti-MPO GN. (**A**) Kidney sections from passive ANCA-MPO GN mice receiving either saline control or ivDNase I treatment stained for NETS as seen by co-localization of MPO (red), H3Cit (green), PAD4 (white) and DAPI (blue) indicated by arrows (**B**) NETS quantitation in glomeruli of ivDNase I treated and control passive ANCA-MPO GN animals. (**C**) Kidney sections labelled for MPO (red), CD45 (green) and DAPI (blue) were assessed for intracellular MPO (co-localizations of CD45 and MPO) and extracellular MPO (non-co-localized CD45 and MPO, arrows). (**D**) Extracellular MPO quantitation in glomeruli of ivDNase I treated and control passive ANCA-MPO GN animals. *P<0.05, ***P<0.001. Data are means ± SEM from 8 mice in the control group and 7 in the treatment group analyzed by Mann-Whitney U. MPO, myeloperoxidase: PAD4, peptidyl arginine deiminase; H3Cit, Citrullinated Histone 3; DAPI, 4’, 6-diamidino-2-phenylindole, Ctrl, Control; DNase I, Deoxyribonuclease I; CD45, pan leukocyte marker C45. Original Magnification 600x

**Figure 8.**
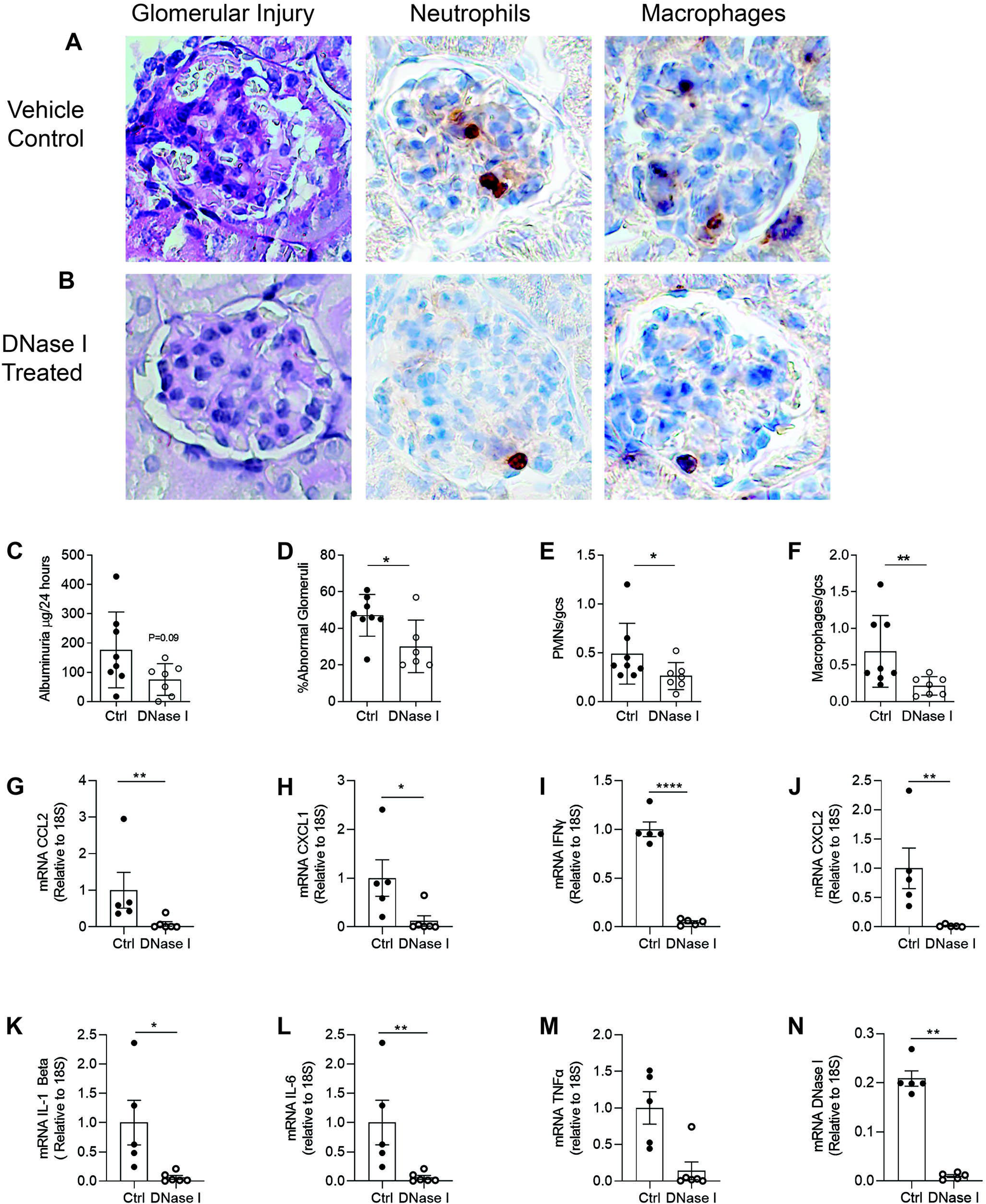
DNase I treatment reduces glomerular injury and inflammatory gene expression in the kidney of mice in an ANCA mediated model of anti-MPO GN. Kidney sections from animals receiving saline-control or DNase I-treatment after passive ANCA-induced anti-MPO- GN were enumerated for evidence of functional glomerular injury as seen by (**A-B**) abnormal glomeruli, polymorphonucleocyte infiltration and macrophage infiltration and (**C**) 24 hour albuminuria levels, (**D-F**) semi-quantification of histological damage and leukocyte infiltration, RNA was extracted from the kidneys of animals receiving saline-control or DNase I-treatment after passive ANCA-induced anti-MPO-GN and assessed by qRT-PCR for (**G**) CCL2, (**H**) CXCL1, (**I**) IFN-γ, (**J**) CXCL2, (**K**) IL-1β, (**L**) IL-6, (**M**) TNFα, (**N**) DNase I. Values are normalized to 18S ribosomal RNA. *P<0.05, ** P<0.01, ***P<0.001, ****P<0.0001. Data are mean± SEM from 8 mice in the control group and 7 mice in the treatment group analyzed by Mann-Whitney U. Ctrl, Control; DNase I, Deoxyribonuclease I. Original Magnification 400x.

**Figure 9.**
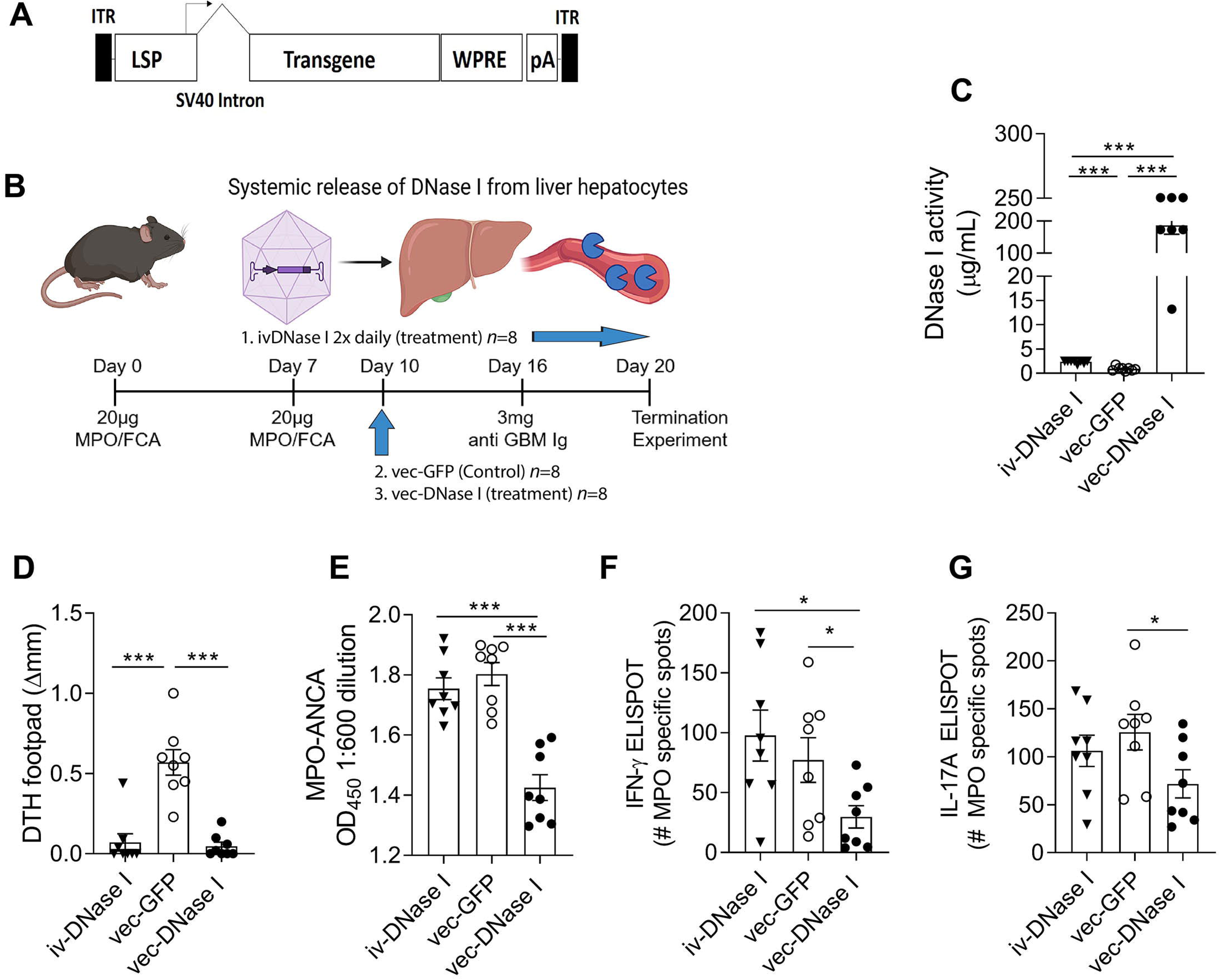
vec-DNase I treatment is superior to exogenously administered DNase I in reducing autoimmunity to MPO in the 20-day ANCA anti-MPO GN model. (**A**) Vector constructs used in the (**B**) the 20-day ANCA model were assessed for (**C**) serum DNase I activity levels. (**D**) After completion of the experiment, mice were assessed for footpad delayed type hypersensitivity response in response to MPO, (**E**) MPO-ANCA levels, and the frequency of anti-MPO reactive cells in the lymph nodes that drained the site of MPO immunization that secreted (**F**) IFN-γ or (**G**) IL-17A. *P<0.05, ** P<0.01, ***P<0.001. Data are mean± SEM from (n=8 mice per group) analyzed by Mann-Whitney U. vec, Adeno associated viral vector; ANCA, anti-neutrophil cytoplasmic antibody; Ctrl, Control; DNA, Deoxyribonucleic acid; DNase I, Deoxyribonuclease I;GFP, green fluorescent protein; H3Cit, Citrullinated Histone 3;MPO, myeloperoxidase.

### Adeno-associated viral vector delivery of DNase I has enhanced therapeutic efficacy over ivDNase I to reduce inflammation in MPO-ANCA GN

A limitation of the therapeutic potential of DNase I that it only has a half-life of < 5 hours. The ivDNase I treatment of the MPO-ANCA required twice daily doses to compensate for short serum half-life of the enzyme. We investigated whether delivery of DNase I could be facilitated using an adeno-associated virus vector to systemically deliver recombinant DNase I (vec-DNase I, Figure 19A). In our previous study, we have shown that the liver provides a continuous super-physiological supply of DNase I in the circulatory system within 3 days of vector injection that stably persists for at least 6 months (34). Using our anti-MPO GN model where disease is mediated by active autoimmunity to MPO, we gave a single dose of vector encoding either DNase I (vec-DNase I) or GFP (vec-GFP) 10 days after the establishment of anti-MPO autoimmunity (Figure 9B). This resulted in a ∼200-fold increase in DNase I activity compared to either ivDNase I delivered twice daily or vec-GFP control treated animals (Figure 9C). vec-DNase I significantly reduced DTH similarly to ivDNase I (Figure 9D). Interestingly, vec-DNase I reduced the generation of MPO-ANCA whereas ivDNase I had no effect on MPO-ANCA levels (Figure 19E). vec-DNase I significantly reduced the number of both IFNγ and IL-17A MPO-specific effector cells in the draining lymph nodes (Figure 9F-G, similar to effects seen with intravenous DNase I administration in the previous sections). Importantly, vec-DNase I also significantly reduced 24-hour albuminuria, whereas ivDNase I administration did not reduce albuminuria (Figure 10A). Further evidence for prevention of glomerular injury by vec-DNase I was seen with reductions in both proportions of glomeruli affected by segmental necrosis, reduction that was more substantial than in iv-DNase I treated mice (Figure 10B) and recruitment of glomerular neutrophils, macrophages and CD4 T cells (Figure 10C-E). To further investigate the role of DNase I in clearing ecDNA we investigated caspase 3 and RIPK3 glomerular protein expression as markers of apoptosis and necroptotic cell death. Vector delivery of DNase I reduced expression of these proteins in comparison to the GFP-control vector group (Figure 10F). Collectively, these data show that gene therapy overcomes the issue of short half-life of DNase I to provide a superior treatment option for anti-MPO GN.

**Figure 10.**
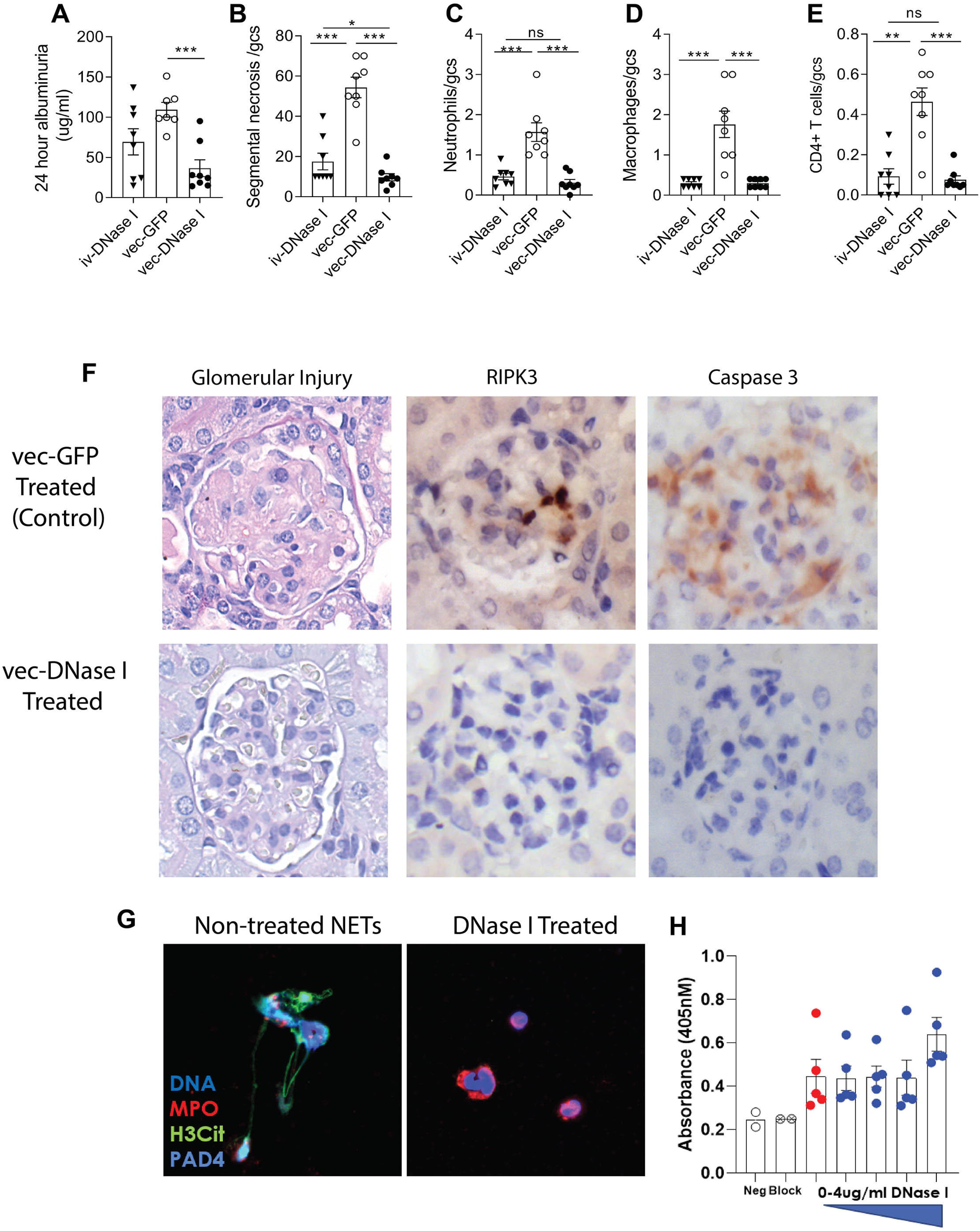
vec-DNase I treatment reduces kidney injury, glomerular leukocytes infiltration, cell death and NETosis. Treatment with vec-DNase I significantly reduces 24-hour albuminuria (**A**) and histological glomerular injury compared to the control GFP vector (**B**). Glomerular leukocyte recruitment is reduced in mice treated with both the exogenous DNase I and vec-DNase I vector when compared to the control AdAV-GFP vector (**C-E**). Representative figures of a reduction in both segmental necrosis and markers of cell death RIPK3 and Caspase 3 are reduced in mice treated with the vec-DNase I vector compared to the control vec-GFP vector (**F**). Neutrophil NET assays stimulated with PMA and treated with DNase I at 1ug/ml demonstrate attenuation of NETs, with DNA in blue, MPO in red, H3Cit in green and PAD4 in grey/blue (**G**). Zymosan phagocytosis assays with neutrophils incubated with DNase I at increased concentrations 0-4μg/ml (blue dots) show that DNase I does not inhibit neutrophil phagocytosis when compared to positive-control (red dots) (**H**). *P<0.05, ** P<0.01, ***P<0.001. Data are means± SEM from 8 mice in each group analyzed by Mann-Whitney U. vec, Adeno associated viral vector;Ctrl, Control; DNA, Deoxyribonucleic acid; DNase I, Deoxyribonuclease I;GFP, green fluorescent protein; H3Cit, Citrullinated Histone 3;MPO, myeloperoxidase; PAD4, Peptidyl arginine deiminase 4, PMA, Phorbal myristate acetate.

### Exogenous DNase I concentrations required to inhibit NETs has no effect on neutrophil phagocytosis

Administration of vec-DNase I can produce more than six months of supraphysiological amounts of DNase I (34). A concern of having a continuous supply of DNase I is that ecDNA released via infectious agents, such as CpG from bacteria, may fail to trigger pattern recognition receptors and associated pro-inflammatory cascades required to eliminate pathogens. To investigate whether using DNase I would compromise neutrophil phagocytosis, and impede host protection, we conducted both NET (Figure 10G) and zymosan phagocytosis assays (Figure 10H) using thioglycolate elicited mouse peritoneal neutrophils and PMA as a stimulant. The lowest concentration of DNase I to prevent NET formation was 1μg/ml (Figure 10G) whereas all concentrations of DNase I between 0-4μg/ml showed no significant differences between phagocytosis controls (red dots) and neutrophils incubated with DNase I (blue dots) (Figure 10 H) indicating that DNase I has no effect on neutrophil phagocytosis at any of the DNase I concentrations. These *in vitro* results with the *in vivo* animal work suggest that DNase I can dampen inflammation caused by NETs whilst still permitting neutrophil phagocytosis by healthy neutrophils.

## Discussion

It is widely accepted that NETs are a major driver of inflammation in MPO-ANCA GN through deposition of MPO, the autoantigen in the disease (16, 19, 28, 29). What is unclear is the contribution of ecDNA in perpetuating inflammation in MPO-ANCA GN. The current study strongly supports a role for ecDNA acting as a pro-inflammatory DAMP mediating injury and autoimmunity in MPO-ANCA GN. We show that ecDNA is a major feature in MPO-ANCA GN patient biopsies and that this is accompanied with a significant diminution of kidney DNase I expression. These observations were mirrored in two animal models of anti-MPO GN that show similar pathophysiology to human disease. We have also shown, that administration of exogenous DNase I in experimental anti-MPO GN diminished NETs, MPO and ecDNA deposition and preserved kidney DNase I expression levels. Moreover, we show for the first time in anti-MPO GN that gene therapy using an adeno-associated viral vector can deliver therapeutic levels of DNase I with just one dose of vector.

Our first aim in this study was to assess kidney biopsies from patients with MPO-ANCA-GN for ecDNA and DNase I expression. MPO-ANCA GN biopsies had significant ecDNA deposits accompanied with a diminution in the protein expression levels of renal DNase I when compared to control patients (minimal change disease). This indicated that DNA may play a significant injurious role in this disease, and that kidney DNase I is depleted in an attempt to clear the ecDNA. NETs are integral to the pathophysiology of ANCA vasculitis, they can be induced by ANCA (plus ANCA independent mechanisms) and are closely associated with the deposition of extracellular MPO within MPO-ANCA GN patient biopsies (16, 29, 35). MPO bound to ecDNA has greater biological activity and is shielded from endogenous inhibitors such as ceruloplasmin (30). Together, this evidence indicates NETs are both a major driver of inflammation and source of autoantigens (16, 29, 35, 36). The capacity of exogenous DNase I to clear not only NET deposited injurious ecDNA but also DNA deposited from all forms of cell death reported in anti-MPO GN (necroptosis, apoptosis and necrosis) was tested in a 20-day murine model of actively induced autoimmune anti-MPO GN, which shows many of the hallmarks of human disease (19, 29, 37). MPO-ANCA GN can be characterized as both a type II antibody mediated disease (due to the effects of ANCA) and a type IV cell-mediated hypersensitivity (due to the Th1 cell mediated aspect of the disease) as per Gell and Coombs classification (38). We and others have shown that deposited extracellular MPO can be recognized by MPO-specific CD4^+^ T cells that in turn initiate cell-mediated glomerular injury characterized by DTH effectors, including the glomerular recruitment of M1 macrophages and the deposition of fibrin, features of GN that are prominent in human and murine anti-MPO GN (39, 40). Additional notable features of the model include generation of MPO-ANCA, patterns of glomerular injury and neutrophil infiltration (focal and segmental necrosis and leukocyte infiltration) and pathological albuminuria (41–44). Administration of DNase I to mice with established anti-MPO GN significantly reduces ecDNA, glomerular NET formation and MPO deposition. DNase I treatment also significantly attenuated MPO-specific dermal DTH and the numbers of glomerular CD4 T cells, macrophages and neutrophils, indicative of DNase I inhibiting type IV cell mediated disease. Importantly, exogenous DNase I treatment also preserved DNase I protein expression levels in the kidney, which is important as depletion of DNase I expression in the organ would render it even more vulnerable to pathological assault induced by uncleared NETs and DNA.

Having established that DNase I is therapeutic even after establishment of anti-MPO autoimmunity in the 20-day cell mediated model of anti MPO GN, we next dissected mechanisms by which DNase I reduces systemic autoimmune responses to MPO. While the mechanisms of DNase I clearance of ecDNA are well understood (27, 45, 46), the mechanisms by which this ecDNA clearance attenuates anti-MPO autoimmunity and GN is unclear. To investigate this phenomenon, we utilized a 10-day model free of complicating GN to dissect out the role of DNase I in preventing systemic autoimmunity to MPO. We revealed that continued daily treatment using exogenous DNase I after animal immunization against MPO using FCA resulted in significant reductions in MPO-specific IFNγ producing cells as well as MPO-induced dermal DTH responses. However exogenous DNase I treatment failed to reduce MPO-ANCA levels. These results confirm a role for DNA in the induction of anti-MPO cellular autoimmunity. The source of this DNA comes from both the accompanying adjuvant, known to contain significant amounts of DNA, and from endogenous neutrophils attracted to sites of MPO immunization whereby they release NETs and DNA (47). Further supporting evidence for a role for DNA activation in MPO-ANCA GN comes from the capacity for DNA subcomponents (e.g. CpG, bacterial DNA components) alone to provide sufficient adjuvant support to allow MPO immunization to induce anti-MPO autoimmunity and GN (48, 49). Furthermore, in the 18-hour model, to dissect out the role DNase I may be playing at the time of antigen presentation, our studies show that DNase I treatment prior to MPO immunization significantly reduces the frequency of DCs in lymph nodes that drain immunization sites and that the DCs that are present show reductions in DC activation markers, including surface costimulatory molecules essential for naïve anti-MPO specific CD4 T cell activation and proliferation. DNase I inhibition of DC activation and lymph node migration supports the view that DNA (in adjuvants or recruited neutrophils) is a potent inducer of MPO autoimmunity.

DNA provides a danger signal to both macrophages and DCs, which induces them to increase their expression of MHC class II expression and co-stimulatory molecules CD40 and CD80/86 (46, 47, 50). The mechanism of DNA activation involves selective DC uptake of the autoantigen MPO by the scavenger receptor DEC-205 followed by binding to the DNA cytosolic receptor TLR9 (49, 51). Therefore, it is likely that DNase I acts by reducing local and/or systemic ecDNA levels thereby circumventing activation of DC and subsequent T and B cell proliferation. MPO-pulsed DCs stimulated with the TLR9 ligand CpG (a component of DNA) stimulates Th1 responses, MPO autoimmunity and promotes kidney injury in experimental anti-MPO GN (49). In contrast, mice receiving *Tlr9*^-/-^ DCs are protected from injury. Given the major ligand of TLR9 is DNA, this provides further supporting evidence that DNA is inflammatory and gives impetus to therapeutically target ecDNA (49). Furthermore, the observation that mice deficient in DNase I or DNase II develop systemic autoimmunity and glomerulonephritis provides additional support for a homeostatic/protective role for DNase I or DNase II (26, 52). Reinforcing the potential of DNase I treatment for acute renal injury are recent studies showing that exogenous DNase I is protective in a rat model of ischemia/reperfusion-induced acute kidney injury and also reduces necroptosis in murine ANCA vasculitis (19, 45).

Having established the kinetics and consequence of early cell-mediated responses in the models, we then turned our attention to defining the role of ecDNA in the production and effects of MPO-ANCA, as this is highly relevant to human disease. ANCA binds to neutrophils inducing their infiltration into glomerular capillaries where they degranulate, produce ROS, undergo NETosis, deposit MPO extracellularly and mediate GN. To test the robustness of our findings in the cell-mediated 20-day model of anti MPO GN we used a second model. We administered DNase I before administering LPS and passively transferring anti-MPO antibodies. In this 6 day model DNase I inhibited anti-MPO antibody-mediated injury by reducing the amount of ecDNA able to activate innate immune inflammatory injury through DAMP pathways.

DNase I treatment of *in vitro* neutrophils has shown that DNase I is associated with a decrease in ROS levels and does not directly inhibit MPO, but degrades the extra-cellular matrix of MPO-associated DNA, which decreases the biological activity of the MPO (53). These modulating effects on neutrophils in the circulation are likely to account for the significant reduction of ANCA-induced neutrophil accumulation in glomeruli. Exogenous DNase I abrogates NET formation, ecDNA deposition and significantly reduces glomerular neutrophil accumulation resulting in significant reductions in pathological injury (assessed by focal segmental necrosis). In this model of MPO-ANCA GN, the effects of DNase I on ANCA-mediated injury can be assessed without any confounding effects of DNase I on anti-MPO autoimmunity, as the transfer of anti-MPO antibodies means that active T and B cell autoimmunity is absent. However, as both these models study acute injury, they do not assess the effects of DNase I over the longer-term development and progression of anti-MPO autoimmunity and ANCA-associated GN.

The current study demonstrates that viral vector delivery of DNase 1 in the 20-day model was able to diminish kidney injury (segmental necrosis and albuminuria). Vec-DNase I reduced the titers of MPO-ANCA, whereas twice daily exogenous DNase I administration did not achieve this outcome. Exogenous delivery of DNase I has also shown to be ineffective at lowering IgA and IgE levels in an animal model of IgA nephropathy (54).It is possible that the sustained and supraphysiological levels of DNase I achieved with vec-DNase I is more effective in removing the effects of ecDNA acting as a DAMP than ivDNase I which has a shorter half-life. This demonstrates for the first time, that vec-DNase I may be used as a biological therapeutic to attenuate the development and/or perpetuation of anti-MPO autoimmunity. As NETs are also a prominent pathophysiological feature that correlate with disease severity and depleted circulatory DNase I in patients with IgA vasculitis, it would be of great interest to see if vector delivery of DNase I has therapeutic benefit in IgA vasculitis (55).

The human protein atlas shows that the kidney has one of the highest mRNA expression levels of DNase I compared to other organs in the body, but the atlas does not show DNase I protein expression within the kidney (56, 57). This is at odds with our results and others that show strong staining for DNase I in kidney tissue from patients with minimal change disease and that enzyme expression is depleted in patients with MPO-ANCA GN. Previous studies in lupus nephritis and membranous nephropathy have also demonstrated DNase I expression both in human and mouse studies (27, 58, 59). In particular, MPO-ANCA GN patients have enhanced loss of DNase I protein within the kidney compared to lupus nephritis patients. This was also observed in our models of anti-MPO GN where OVA-immunized mice (without autoimmunity to MPO) had significantly more DNase I protein expression than untreated mice and exogenous DNase I partially restored this level in the kidney.

DNase I has been shown to be critical in preventing chronic neutrophilia. *Dnase1^-/-^ Dnase1l3 ^-/-^* mice, deficient in both proteins have intravascular occlusions, with NETs entrapping platelet and erythrocytes (60). DNase 1L3 is expressed in immune cells, whereas DNase I is secreted by non-hematopoietic tissue. Restoring either of the DNase enzymes in double knock-out mice prevented neutrophil accumulation, NET formation and premature death. This adds weight to the argument that constant systemic release of DNase I is required to clear neutrophil derived DNA.

There has been considerable interest for many decades in clinically translating recombinant DNase I in multiple different disease contexts. In the pre-antibiotic era, the enzyme was used topically to clear and sterilize infected wounds and relieve meningitis (61). Currently, it is administered in nebulized form to the airways of cystic fibrosis patients to degrade nuclear material released from dead neutrophils, thereby reducing viscosity of respiratory secretions (62). Intravenous DNase I has also been used in clinical trials of systemic lupus erythematosus where it was safe and well-tolerated, although ineffective in disease control possibly because of its relatively short half-life in serum (<5 hours)(63). Our study has shown gene delivery provides a solution to this issue and creates an option that is therapeutically superior to intravenous enzyme delivery. Adeno-associated virus vectors have a strong safety profile and permit long-term transgene expression in post-mitotic cells (in the liver) making them ideally suited to treat chronic and remitting diseases such as MPO-ANCA GN, which typically requires ongoing treatment that may be prolonged over years (64–66). Additionally, use of a vector that targets hepatocytes to release DNase I systemically rather than targeting the kidney directly circumvents the limitations of successful gene transfer of damaged kidney cells. As MPO-ANCA GN patients have significantly damaged kidneys, in particular endothelial cells and podocytes, therefore tissue pathology may be a barrier to successful gene transfer. Systemic release of DNase I via the liver hepatocytes will not only target ecDNA present in the kidney but will have further therapeutic benefit by also digesting DNA being released from cell death within the circulation. Further development of the technology to pharmacologically regulate transgene expression would offer the remarkable prospect of switching therapy “on and off” to regulate vector dosing of DNase I, thus allowing for therapy during active disease that could be discontinued during remission and re-activated in relapsing disease (67). The therapeutic efficacy of vector-delivered DNase I in a murine model that emulates human MPO-ANCA GN supports the possibility that this gene therapy may be a future approach in treating clinical disease in people with ANCA-associated vasculitis.

## Materials and Methods

### Sex as a biological variable

As MPO-ANCA GN affects both male and females equally, both sexes were included in human and mouse studies.

### Patient Cohort

Twenty-nine renal biopsies from consented patients with diagnosed MPO-ANCA GN, and 6 renal biopsies from patients with Minimal Change Disease (MCD), a non-proliferative form of kidney disease with normal light microscopy were used in this study (clinical data in Supplementary Table 1). Biopsies were collected from consented patients with ethics approval over a period from 2001-2016 at Monash Health, Clayton, Victoria, Australia. The biopsies from MPO-ANCA GN patients were collected at the patient’s first presentation.

### Mice

C57BL/6 mice were bred at Monash Medical Centre Animal Facilities and used at 8-10 weeks of age. *Fcgr2b*^-/-^ mice (C57BL/6J background: B6; 129S-Fcgr2b^tm1Ttk^/J) were purchased from the Jackson Laboratory (Bar Harbor, ME) and bred at Monash Medical Centre Animal Facilities, Monash University, Australia. The phenotype of *Fcgr2b*^-/-^ mice was confirmed by flow cytometry with anti-B220 and anti-CD16/CD32Ab (BD Biosciences, North Ryde, NSW, Australia), as previously described (68). Mice aged 8-10 weeks were used for experiments under specific pathogen-free conditions.

### Experimental Design

An established model of active anti-MPO autoimmunity was used to assess anti-MPO cell mediated glomerular injury (41–44). WT mice (male and female) were immunized intraperitoneally with 20 µg recombinant murine (rm) MPO (69) in Freund’s complete adjuvant (FCA; Sigma-Aldrich) and boosted subcutaneously with 10 µg rmMPO in Freund’s incomplete adjuvant (FIA; Sigma-Aldrich) on day 7. Disease was initiated (‘triggered’) by intravenous injection of 3mg of anti-GBM globulin on day 16 (administered as 1.5mg, 2 hours apart), 4 hours prior to treatment. Anti-MPO autoimmunity and GN was assessed 4 days later (day 20). As a control for the immunization process and the anti-GBM globulin one group of control mice received an irrelevant antigen, OVA instead of MPO. Treated mice with anti-MPO GN received DNase I 10mg/kg (Pulmozyme, Roche, Basel Switzerland) twice daily in 300 μl of normal saline intravenously, commencing 4 hours after the last anti-GBM globulin to ensure recruited neutrophils had deposited MPO to initiate disease, and twice daily until termination of the experiment on day 20. Non-treated control mice were immunized with MPO and injected with anti-GBM globulin, then received vehicle injections of saline alone. For experiments using AAV8-DNase I both the control GFP vector and DNase I vector were given on day 10 (intraperitoneal injection of 1 ×10^11^ vector genomes [vg per mouse]) and compared to intravenous delivery of recombinant DNase I (Pulmozyme, 10mg/kg) administered twice daily (12 hours apart) from day 16 after triggering of disease with the anti-GBM antibody until termination of the experiment on day 20.

To assess MPO-ANCA (humoral) induced glomerular injury, anti-MPO IgG generated by immunizing *Mpo-/-* mice as previously described (70) was passively transferred into *Fcgr2b^-/-^* mice, which are more susceptible to anti-MPO GN (71). Mice were primed with intraperitoneal LPS (0.5µg/g) 2 hours prior to intravenous transfer of anti-MPO IgG, which activates and recruits neutrophils to glomeruli. DNase I was given twice daily intravenously on day −1 and twice daily until the end of experiments on day 6.

### Adeno-associated viral vector production, DNase I activity assay and quantification of liver vector copy number

AAV vectors encoding GFP or and DNase I packaged into AAV serotype 8 (AAV8) capsids were purified and titrated as previously (34). Vectors were injected into the intraperitoneal cavity at 1 x 10^11^ vector genomes (vg) per animal. At the end of the 20-day anti MPO-GN model livers were removed and examined for vector copy number as previously published (34). Vector copy number was determined using Platinum Taq DNA polymerase (Invitro-gen) according to the manufacturer’s instructions, using primers specific for the WPRE region on the vector and normalized to GAPDH. DNase I activity from plasma samples were determined using radial enzyme diffusion assays according to our previously published protocol (31, 34). The halo diameter around each well of digested DNA was measured and compared to DNase I standards.

### Extracellular DNA, Neutrophil extracellular Traps and extracellular MPO detection

To measure extracellular DNA, 3 µm formalin-fixed, paraffin-embedded (FFPE) kidney sections were cleared in xylene, rehydrated in graded alcohols, and immersed in antigen retrieval solution (10mM Tris, 1mM EDTA, pH 9.0) and boiled in a pressure cooker for 10 minutes, as described previously (31). Extracellular DNA was assessed by using colocalization studies using 4′,6-diamidino-2-phenylindole (DAPI), (D1306, Thermofisher, Waltham, MA, USA) as a marker for dsDNA (DAPI is nucleic acid specific, which preferentially binds to the AT base pairs in the mirror groove of DNA) and β-actin as a structural component of all cells to delineate glomerular and cell structures (31). Sections were blocked in 10% chicken sera in 5% bovine serum albumin (BSA)/ phosphate buffered saline solution (PBS) for 30 minutes and stained with a rabbit anti-mouse β-actin antibody (ab8227, ABCAM, Cambridge, UK) at 1µg/ml in 1%BSA/PBS, as a cell marker (and to identify glomerular regions of tissue for measurement) overnight at 4°C. For secondary detection, a chicken anti-rabbit Alexa Fluor 488 antibody at 1:200 was used for 40 minutes at RT (Molecular Probes, Thermofisher). Slides were quenched in Sudan Black B as outlined below and mounted with DAPI prolong gold mounting media to detect DNA. NETs and extracellular MPO were detected as published previously (18). Briefly, sections (3 μm) of FFPE tissue specimens were mounted on superfrost plus slides (Menzel, Braunschweig, Germany), dewaxed, rehydrated, and pre-treated with antigen retrieval solution Tris-EDTA pH 9 (10mM TRIS, 1mM EDTA) in a pressure cooker for 10 min, blocked (30 min) in either 10% chicken sera in 5% BSA/PBS (immunofluorescence) and probed with goat anti-human MPO (detects mouse MPO, AF3667 R&D Systems) rat anti-mouse CD45 (all leukocytes, to define extracellular MPO, Becton Dickinson, Franklin Lakes, New Jersey, USA), and mouse anti-human PAD4 (ab128086, ABCAM), rabbit anti-human H3Cit (ab5103, ABCAM), and goat anti-human MPO (AF3667, R&D systems) in 1% bovine serum to detect NETs, diluted in albumin/phosphate-buffered saline for 16 h (4°C). Secondary detection was with either Alexa Fluor 594-conjugated chicken anti-goat IgG or Alexa Fluor 488-conjugated chicken anti-rabbit IgG, or donkey anti-mouse 647 IgG (all from Molecular Probes, Thermofisher 1:200, 40 min, room temperature). To quench tissue auto fluorescence, slides were incubated with Sudan Black B (Sigma-Aldrich St Louis, MO, 0.1% in 70% ethanol, 30 min), washed in phosphate-buffered saline, and cover slipped in DAPI prolong gold (Molecular Probes). Fluorescent images were acquired using a NIKON C1 confocal laser scanning head attached to Nikon Ti-E inverted microscope (Coherent Scientific, SA, Australia); 405, 488, and 561 nm 647nm lasers were used to specifically excite DAPI, Alexa 488, Alexa 594, and Alexa 647. Single plane 512×512×12 bit images were captured in a line-sequential manner (4 line averaging) using a 20×, 40×, or 60× objective.

### Assessment of ecDNA, extracellular MPO and DNase I

To assess ecDNA in human renal biopsies and murine kidneys, a published method using “analyze particle” in image J was used and circularity set to include nuclear staining, the nuclear stain was thresholded, marked and excluded from analysis using the watershed function in Image J, leaving only the extranuclear stain to be measured (NIH, Bethesda, MD) (31). The average integrated intensity/density was used and expressed in arbitrary units per glomerular cross section. Extracellular MPO was measured by a previously published method using a macro in Image J analysis software (NIH, Bethesda, MD)(29). Intracellular (leukocyte-associated) MPO was defined being associated with CD45 (CD45+MPO+ cells). Extracellular MPO was defined and measured as MPO+CD45-staining. The macro evaluated both the area and intensity and expressed the results as arbitrary units (AU).

### Assessment of renal human and mouse DNase I

DNase I immunohistochemistry was performed on 3 μm thick, formalin-fixed, paraffin-embedded human renal biopsies from MPO-ANCA-associated vasculitis patients and MCD patients, and murine models of T cell mediated anti-MPO GN and GN induced by passive transfer of anti-MPO antibodies. Sections were cleared in Histosol, rehydrated in graded alcohols and then blocked with 10% Horse serum in 5% BSA/PBS for 30 minutes at RT and incubated with a mouse anti-human DNase I antibody (1/100, Santa Cruz, overnight at 4 degrees). Sections were washed, and endogenous peroxidase blocked with 1% Hydrogen Peroxide (H_2_0_2_) in methanol for 20 minutes, followed by blocking with avidin and biotin using a commercially available blocking kit per manufacturer’s instructions (Vector Laboratories, Newark, CA, USA). Secondary antibody detection was performed with a horse anti-mouse biotinylated antibody at 1:60 (Vector Laboratories) for 40 minutes at RT, and detected with an avidin biotin complex conjugated to HRP (ABC-HRP) for 40 minutes (Vector Laboratories), then with 3,3′-diaminobenzidine (DAB brown, SIGMA), dehydrated, cleared in Histosol and mounted in Vector Shield (Vector Laboratories) permanent mounting media. Five low power (20x objective lens) fields of view of the tubulointerstitium were analyzed and graded 1-3 according to the amount of stain present within tubulointerstitial epithelial cells in human renal biopsies, and 10 interstitial fields of view were analyzed in mice.

### Assessment of renal injury and leukocyte infiltration

Histologic assessment of renal injury was performed on 3 μm thick, FFPE fixed, periodic acid-Schiff-stained kidney sections. At least 50 consecutive glomeruli/mouse were examined and results expressed as percentage of abnormal glomeruli exhibiting either; crescent formation, segmental necrosis, sclerosis or infiltrating immune cells and expressed per glomerular cross-section (gcs). Glomerular CD4+ T cells, macrophages, and neutrophils were assessed by an immunoperoxidase-staining technique on 6 μm thick, periodate lysine paraformaldehyde (PLP) fixed, OCT frozen kidney sections. The primary antibodies used were GK1.5 for CD4+ T cells (anti-mouse CD4+; American Type Culture Collection, Manassas, VA), FA/11 for macrophages (anti-mouse CD68 from Dr. Gordon L. Koch, Cambridge, England), and RB6–8CS for neutrophils (anti-GR-1; DNAX, Palo Alto, CA). At least 30 glomeruli were assessed and results expressed as cells/gcs (c/gcs).

### Systemic Immune Responses to MPO

ELISA was used to detect circulating serum anti-MPO IgG titers using 100µl/well, 1µg/ml murine MPO and horseradish peroxidase conjugated sheep anti-mouse IgG (1:1000; Amersham Biosciences, Rydalmere, Australia). IFN-γ and IL-17A production was assessed by ELISPOT (BD Biosciences, Mouse IFN-γ ELISPOT kit and Mouse IL-17A ELISPOT kit) with draining LN cells seeded at 5×10^5^ cells/well re-stimulated with 10 μg/ml of rmMPO for 18 hours. IFN-γ and IL-17A producing cells were enumerated with an automated ELISPOT reader system (TECAN). To assess MPO-specific dermal delayed type hypersensitivity (DTH), mice were challenged by intradermal injection of 10 μg murine MPO in 30μl saline in the right hind footpad (the contralateral footpad received saline). DTH was quantified 24 hours later by measuring the difference between footpad thicknesses (Δmm) using a micrometer.

### Flow cytometry and intracellular staining

DCs were identified as CD11c^hi^ cells on isolated draining LN cells by flow cytometry. LN cells were stained for 30 minutes at 4°C with the following directly conjugated antibodies: hamster anti-mouse CD11c PE (HL3), hamster anti-mouse CD11c PerCP/Cy5.5 (N418, BioLegend), rat anti-mouse CD40 FITC (3/32, BD Biosciences), rat anti-mouse MHC-II APC/Cy7 (M5/114.15.2, BioLegend), rat anti-mouse ICOS Ligand PE (HK5.3, Biolegend), mouse anti-mouse OX40L PE (8F4, BioLegend), hamster anti-mouse CD80 FITC (16-10A1, eBioscience), rat anti-mouse CD86 APC/Cy7 (GL-1, BioLegend). CD4 (GK1.5, BD), CD69 (H1.2F3, BD). For analysis of MPO specific Treg responses *ex vivo*, 5 x 10^5^ LN cells were cultured for 72 hours with 10µg/ml MPO. CTV, as per manufacturer’s instructions was added at 0 hours and cells stained using anti-Foxp3 (FJK-16s, EBioscience) and Foxp3 fixperm kit (Ebioscience). Cells were analyzed on the Beckman Coulter Navios platform and data analyzed using FlowJo software (TreeStar, Palo Alto, CA).

### NET and Zymosan phagocytosis Assays

For the *in vitro* NET assay and zymosan phagocytosis assay *n*=5 C57BL/6J male mice were injected with aged 4% thioglycolate intraperitoneally (500μl). Neutrophils (2×10^5^) were plated out in 24 well plates for immunostaining, then pre-incubated with DNase I at varying concentrations from 0-4ug/ml DNase I for 30 minutes prior to adding PMA (40µg/ml, SIGMA) for 3 hours at 37°C to induce NET formation in serum free RPMI (SIGMA) media [to avoid blocking of NET production with albumin (72)]. Cells were fixed in paraformaldehyde (2%) supplemented with periodate and lysine overnight at 4°C. Cells were washed, permeabilized, blocked in 10% chicken sera and then stained for markers of NETs, goat anti human/mouse myeloperoxidase (anti-MPO, AF3667, R&D Systems, Minneapolis, MN), rabbit anti human/mouse citrullinated histone 3 (H3Cit, Ab5103, Abcam), and mouse anti human/mouse peptidyl arginine deiminase 4 (PAD4; Ab128086, Abcam), and detected with secondary chicken anti-rabbit AF488 (Thermofisher), chicken and mouse AF647 (Thermofisher), and chicken anti-goat AF594 (Thermofisher) and mounted in DAPI ProLong Gold (Molecular Probes, Thermo Fisher Scientific). NETs were visualized by confocal fluorescence microscopy with a Nikon Ti-E inverted microscope (Nikon Instruments, Melville, NY). Then, 405-, 488-, 561-, and 647-nm lasers were used to specifically excite DAPI, Alexa Fluor 488, Alexa Fluor 594, and Alexa Fluor 647.

For the zymosan phagocytosis assays neutrophils were plated out at 5×10^5^ into a 96 well plate overnight at 37°C treated with 0-4µg/ml DNase I 30 minutes prior to the addition of PMA at 40µg/ml for 3 hours. The zymosan phagocytosis assay was then performed using a standard kit ( ab211156 Abcam), visualized using the kit detection reagent and read at an absorbance plate reader at OD 405nm (Tecan).

### Real-Time PCR for gene expression

Kidneys were stored at minus 80 degrees until time of RNA extraction. RNA extraction was performed via a standard protocol using trizol, a High capacity cDNA reverse Transcription Kit (Applied Biosystems, Foster City, CA) was used to generate cDNA. Taqman Universal PCR Master Mix and Taqman Gene expressions assays were performed as per the manufactures instructions and normalized to the house keeping gene *18S* (Applied Biosystems, Gene specificities for (*Ccl2, Cxcl1, Cxcl2, Ifng, Il1b Il6, Tnf* and *DnaseI*) and read on the 7900HT Fast Real-Time PCR System (Applied Biosystems).

### Statistics

Results are expressed as the mean ± SEM. Kruskal-Wallis one-way analysis of variance was used for comparisons between 3 groups, a Dunn post hoc test was performed for multiple comparisons of groups. Mann–Whitney t-test was used for non-parametric data and paired t-test for comparison in ex vivo co-culture experiments. All data were analyzed with GraphPad Version 6 (GraphPad Prism; GraphPad Software Inc., San Diego, CA). Differences were considered to be statistically significant if *P*<0.05.

### Study Approval

Study approval for use of patient kidney biopsy material and use of clinical data was approved by the Monash Health Ethics Committee (application number 08216B). Written informed consent was received prior to participation. Study approval for the animal work within the study was approved by Monash University’s animal ethics committee approved application numbers; MMCB2019/35, MMCB2014/23, and MMCB2017/43.

## Author contributions

K.M.O., G.J.L., P.Y.G, A.R.K., and I.A.E., conceptualized the study, K.M.O., A.C.L., P.Y.G., J.D., M.A.A., M.M., D.K.Y.C., G.J.L., conducted the experiments, K.M.O., P.Y.G., J.D., M.A.A., M.M., G.J.L., analyzed the data, K.M.O., wrote the first draft, made the figures, and K.M.O., G.J.L., P.Y.G., I.A.E., and A.R.K., reviewed and edited the paper, K.M.O., G.J.L. and P.Y.G., acquired the funding, all authors approved the final version of the manuscript.

## Acknowledgments

We acknowledge funding support from the National Health and Medical Research Council of Australia for a project grant (ID 1146892), an Ideas Grant (ID 2020579) and the Rebecca L Cooper Foundation (PG2019449) for a fellowship; Monash Micro Imaging for the provision of the Nikon, Confocal Microscope, Monash Health Translational Precinct Flowcore for flow cytometry, the Monash Health and Translational Genomics Platform for RT-PCR machine access and the Australian Government for providing a Research Training Program Scholarship.

