## Supplementary Table and Figure for "Treatment of anti-myeloperoxidase glomerulonephritis using recombinant deoxyribonuclease I is enhanced by adeno-associated virus gene therapy"

**Table 1. Clinical Data from patients with ANCA associated Vasculitis**

| ***Patient Characteristics*** | **MPO-ANCA-associated vasculitis** | **MCD** |
|---|---|---|
| **Patient number** | 29 | 6 |
| **Age** | 63.5 (±3) | |
| **Sex F/M** | 15/14 | 4/2 |
| **Number of glomeruli** | 18 | 19 |
| ***Laboratory values*** | | |
| **MPO-ANCA titre (U/ml)** | 149.58 (±23) | n/a |
| **eGFR (ml/min/1.73 $m^2$)** | 34.77 (±8) | 100.1 (±6) |
| **ESR (mm/h)** | 70 (±8) | n/a |
| **CRP (mmol/l)** | 66.4 (±25) | n/a |
| **Urinary red blood cells (cells/HPF)** | 552 (±15) | n/a |

*Abbreviations: ANCA, anti-neutrophil cytoplasmic antibody; CRP, C reactive protein; eGFR, estimated glomerular filtration rate; ESR, erythrocyte sedimentation rate; MCD, minimal change disease MPO, myeloperoxidase*

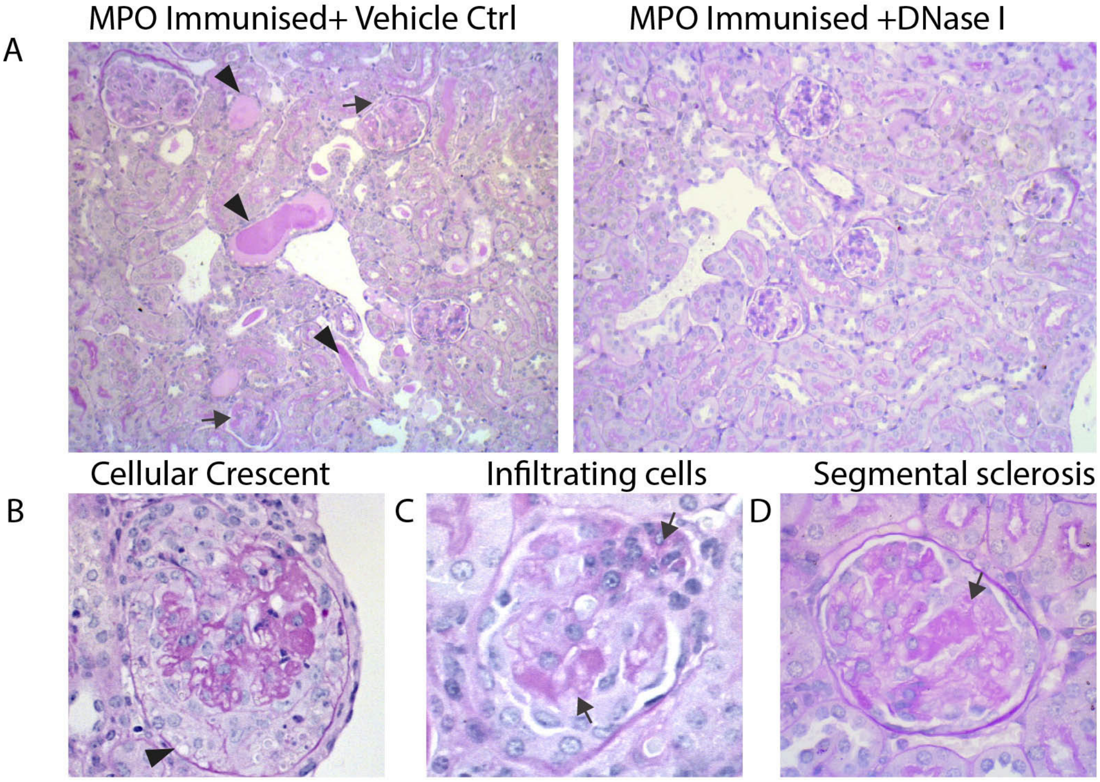


**Supplementary Figure 1. Low powered images demonstrating extent of histological damage between the vehicle control group and treatment groups and examples of histological injury observed.** (**A**) Low powered Image of the tubulointerstitium showing multiple tubular casts (arrow heads) and damaged glomeruli (arrows) compared to DNase I treated kidney with no tubular casts and minimal histological changes to glomeruli. (**B**) Example of cellular crescent, arrow pointing to crescent within Bowman's capsule(**C**) Infiltrating cells into Bowman's space and area of segmental necrosis, arrows(**D**) examples of segemental sclerosis/necrosis. Abbreviations: MPO, myeloperoxidase, DNase I, deoxyribonuclease I, Ctrl. Control. Original magnification for (A-B) 200x, original magnification for (B-D) 400x.
